# Fragmentation reduces vesper bat abundance and evenness in a southern Appalachian forest

**DOI:** 10.1101/2025.11.08.687270

**Authors:** Ethan J. Ramey, Thaddeus R. McRae

## Abstract

Habitat fragmentation alters biodiversity through changes in habitat patch size, structure, and increasing distance between patches. However, if species regularly move across distances greater than those between patches, the effect of distance may approach zero. More studies of highly mobile species such as vesper bats (*Vespertilionidae*) are needed to clarify how such species respond to fragmentation. This study examines the impact of forest fragmentation on vesper bat community diversity in the Southeastern United States. Using mobile acoustic surveys, bats were monitored along forest transects in the Cherokee National Forest and in surrounding farmlands containing widely spaced forest fragments. The same nine vesper bat species were recorded and identified in both habitat types. Results indicate a 43% decline in vesper bat abundance in the farmland when compared to intact forest, with eight species having a higher abundance within the intact forest. Notably, Seminole bats, (*Lasiurus seminolus*) maintained similar abundance across habitats, suggesting species-specific responses to fragmentation. Despite identical species richness, diversity indices such as the Shannon diversity index and *β_Shannon_* identified lower biodiversity in the farmland when compared to the forest. Our results indicate that despite being highly mobile, vesper bats exhibit a largely negative response to habitat fragmentation and loss. This could be due to the change of the core to edge habitat ratio, emphasizing the importance of conserving core forest habitat to sustain bat biodiversity. Observed species-specific responses to habitat fragmentation indicate a need for further research to elucidate the complexities of managing bat communities in human-modified ecosystems.

## Introduction

Habitat fragmentation has consistently been shown to have profound effects on species diversity and community composition (Fahrig 2017; Kupfer et al. 2006; Gorresen and Willig 2004; Laforge et al. 2022). Although habitat fragmentation clearly affects biodiversity, the nature and magnitude of its effects are complex and differ across study systems and spatial scales (Chetcuti et al. 2020). In a review of the effects of fragmentation, Fahrig (2017) found that 76% of reported statistically significant responses to habitat fragmentation were positive in regard to biodiversity; in contrast, many studies (Fletcher 2005; Haddad *et al*. 2015; Atkinson and Shorrocks 1981; Rybicki, Abrego, and Ovaskainen 2020; Püttker et al. 2013) have noted negative effects on diversity in fragmented habitat. It is well agreed upon that core species are better suited to larger fragments (Ewers and Didham 2007; Haddad et al 2015; Szitár et al 2023). Consequently, in some cases, habitat fragmentation causes extinction of core-species populations by decreasing core habitat and increasing forest edge habitat and edge effects (Ewers and Didham 2007).

### Habitat fragmentation affects biodiversity in diverse ways

There are at least four distinct ways that habitat fragmentation may have a positive or negative effect on biodiversity; changing ratio of edge to core habitat, reduced patch size, increased patch isolation, and changes in habitat type with patches (e.g., forest structure changes).

#### Fragmentation affects populations by changing ratio of edge versus core habitat

While moderate habitat fragmentation may increase the amount of edge habitat and thus the abundance of edge species, additional fragmentation will eventually decrease edge habitat as well as core habitat, due to the associated habitat loss (e.g. removal of forest canopy). This complex response to fragmentation is in part due to differences in response at the species level between edge species and core species. Fragmentation creates more edge habitat, which increases suitability for species who thrive at the forest’s edge. In Eastern North American systems, species such as white-tailed deer (*Odocoileus virginianus*), (Dechen Quinn et al. 2013), and various indigenous songbirds (Parker et al. 2005) have been shown to benefit from increased edge habitat in fragmented ecosystems. Conversely, the loss of core habitat as fragmentation continues has resulted in negative effects on some species. While generalists may well adapt to fragmented habitat, specialist species tend to have a difficult time adapting to fragmentation due to their specific habitat requirements or limited dispersal abilities, or both (Henle 2004). Wood thrushes, a forest interior species, experienced a decline in abundance between 59-79% in forest fragments that are experiencing nearby development (Heide et al. 2023). Glanville fritillary butterflies (*Militae cinxia)* were divided into smaller populations by habitat fragmentation, which forced inbreeding; ultimately threatening local extinction (Hanski et al. 2011). A study in the Brazilian Atlantic Forest found that three out of four forest specialist rodent species disappeared when fragmentation reduced forest coverage to 10% (Pardini et al. 2010). Species that are highly dependent on forest canopy cover experience reduced γ diversity in response to habitat fragmentation (Chetcuti et al. 2020). Although fragmentation inherently co-occurs with habitat loss, because some amount of the original habitat type is always removed to create fragments, the γ diversity reduction observed by Chetcuti et al. (2020) was independent of any effects of habitat loss itself. Thus, fragmentation itself, when all habitat patches remain reachable, will equate to habitat loss for core habitat species and habitat gain for edge habitat species, thus shifting the ratio of edge to core. However, as fragmentation increases, the overall ratio of original habitat (edge plus core) to matrix decreases. Eventually a tipping point is reached where total area of edge habitat as well as core habitat is reduced by further fragmentation, thus threatening both core and edge species.

#### Biodiversity decreases from fragmentation reducing patch size

Negative effects on biodiversity could arise from high degrees of fragmentation resulting in habitat patches that are too small to sustain a healthy community. Haddad *et al*. (2015) reported that habitat fragmentation can reduce biodiversity by 13-75%. A survey of protected areas (PAs) across the world found that 34% of these PAs experienced habitat fragmentation, 19% of PAs experienced habitat loss, and 10 of these contained endangered species that are critically threatened by habitat loss and fragmentation (Yuan et al. 2024). Further, a study of 147 European grassland fragments observed significant immediate and time-delayed declines of biodiversity in response to fragmentation (Krauss et al. 2010).

#### Biodiversity decreases from fragmentation increasing distance between patches

Certain species’ dispersal ability may be insufficient to cross the matrix between fragments of habitat (Lu, *et al*. 2012). Püttker et al. (2013) observed a decrease in immigration of two habitat specialist rodent species in smaller, fragmented patches compared to those in habitats with more canopy coverage; suggesting that gene flow between certain populations is threatened by habitat fragmentation. Further, patches of fragmented habitat that are more widely spaced may prevent less-mobile species from moving between patches, essentially rendering those relatively remote patches as lost habitat.

Additionally, edge effects from fragmentation could reduce biodiversity (Fahrig, 2003; Fletcher, 2005). 70% of the world’s remaining forest is within 1 kilometer of the forest’s edge (Haddad et al. 2015). Increasing the gap size between these patches could decrease the reproductive rate of certain species (Fahrig 2002). Thus, the importance of studying the implications of habitat fragmentation and edge effects cannot be overstated.

#### Biodiversity can increase from fragmentation increasing edge habitat and diversifying forest structure

Positive effects of fragmentation on biodiversity have also been observed. Some potential reasons for these positive effects include positive edge effects as mentioned above (Tanner 2005; Wordley et al. 2018), the habitat gradient created by fragmentation (Kupfer *et al*. 2006), and that forest structure and composition may diversify due to fragmentation, thus increasing diversity of foraging, resting, and breeding resources for wildlife (White et al. 2022). Since habitat fragmentation can also increase habitat diversity, species may respond positively to the creation of corridors that connect habitat patches (matrices), encouraging dispersal between these patches (Dunning *et al*. 1992) and positively affecting biodiversity (Tscharnke *et al*. 2002).

### Mobile species may experience fragmentation as net habitat loss versus island effects

Although habitat fragmentation can have diverse effects on population sizes and community diversity, as fragmentation becomes extreme, it consistently creates large distances between habitat patches. This distance is likely to have varied effects on populations depending on the ease with which each species can cross the matrix between patches. If a species is highly mobile, so that most patches are readily accessible, then this fragmentation may be experienced primarily as habitat loss, where the distance between patches has less effect than the total habitat area lost. Any such community of highly mobile species that experiences fragmentation primarily as net habitat loss would show consistent decrease in abundance in fragmented habitat, with minimal change to relative abundance of each species. Vesper bats, (*Vespertilionidae*) are one such group of highly mobile species, reaching speeds that range from 13.7 to 43.2 km/h (Cueva Salcedo et al. 1995) as they forage for flying insects at night, making them an effective model to observe how highly mobile species respond to habitat fragmentation. In addition, understanding vesper bat response to fragmentation is key to understanding temperate forest ecology as bats play important roles in ecosystem function, such as preying upon nocturnal flying insects, and reducing top-down pressure on plants (Federico, *et al*. 2008; Beilke and O’Keefe 2022). Given their importance to ecosystem health and the fact that they are facing multiple threats (Frick, Kingston, and Flanders 2020; Hoyt *et al*. 2021), biologists must understand the effects of habitat fragmentation on vesper bat communities in order to make informed decisions about forest management and the conservation of these keystone species (Meyer, Struebig and Willig 2015).

While vesper bats are primarily insectivorous, much of the research surveying bat communities in the context of habitat fragmentation has been done with phyllostomid bats (*Phyllostomidae*), a tropical family of bats with diverse diets with various species specializing on nectar, pollen, or fruit in addition to some species that also prey on animals (Bernard and Fenton 2007; Meyer, Struebig and Willig 2015; Rocha, Ovaskainen, López-Baucells *et al. 2018).* This tropical bat research primarily investigated effects of fragmentation such as foraging behavior (Rodríguez-San Pedro and Simonetti 2015), and ability to disperse across habitat matrices (Fahrig 2007).

While extensive surveys of insectivorous vesper bat community composition have been conducted in the Eastern US, there is still a need for studies designed to explicitly examine the effects of fragmentation by comparing neighboring bat communities in intact versus fragmented forest habitat. To our knowledge, a study comparing Southeastern United States temperate forest bat communities between continuous and fragmented habitat communities like ours has not been published. We set out to compare the presence and relative abundance of vesper bat species between a forest with continuous tree canopy and highly fragmented forest within an agricultural matrix; thereby testing whether this community of highly mobile species will respond with a similar drop in abundance across all species or will show changes in relative abundance and community composition.

## Methods

### Study location

To better understand how vesper bat communities in temperate forest ecosystems are affected by habitat fragmentation we performed mobile acoustic surveys of bat communities near Benton, Tennessee in the Cherokee National Forest and in farmland with fragmented forest patches. We surveyed bats in summer 2024 along road-based transects of similar length in each habitat. Road transects were chosen based on several factors: first, our transects had to be low traffic roads to minimize effects of vehicles on bat behavior and detections, and to enable safe data collection, which requires driving at low speeds. Second, these roads had to be similar in length to ensure equal sampling effort. Third, the two transects had to be as close in proximity as possible, limiting any regional differences between transects.

Our forest transect (Fig. 1) was 25.75 kilometers long, with continuous forest canopy broken only by small clearings around developed areas and roads. There were a variety of surface water features along this transect, we identified three subtransects based on proximity to streams and rivers of various sizes. Forest subtransect A had little to no proximate surface water and was the longest section of the total transect. Forest subtransect B was riparian, directly against the Ocoee River. Finally, forest subtransect C was intermittently surrounded by small, first and second-order streams. While the lengths of the three sections were unequal due to the nature of the available roads and waterways, this division enabled us to test the effects of waterways on bat communities.

**Fig. 1.**
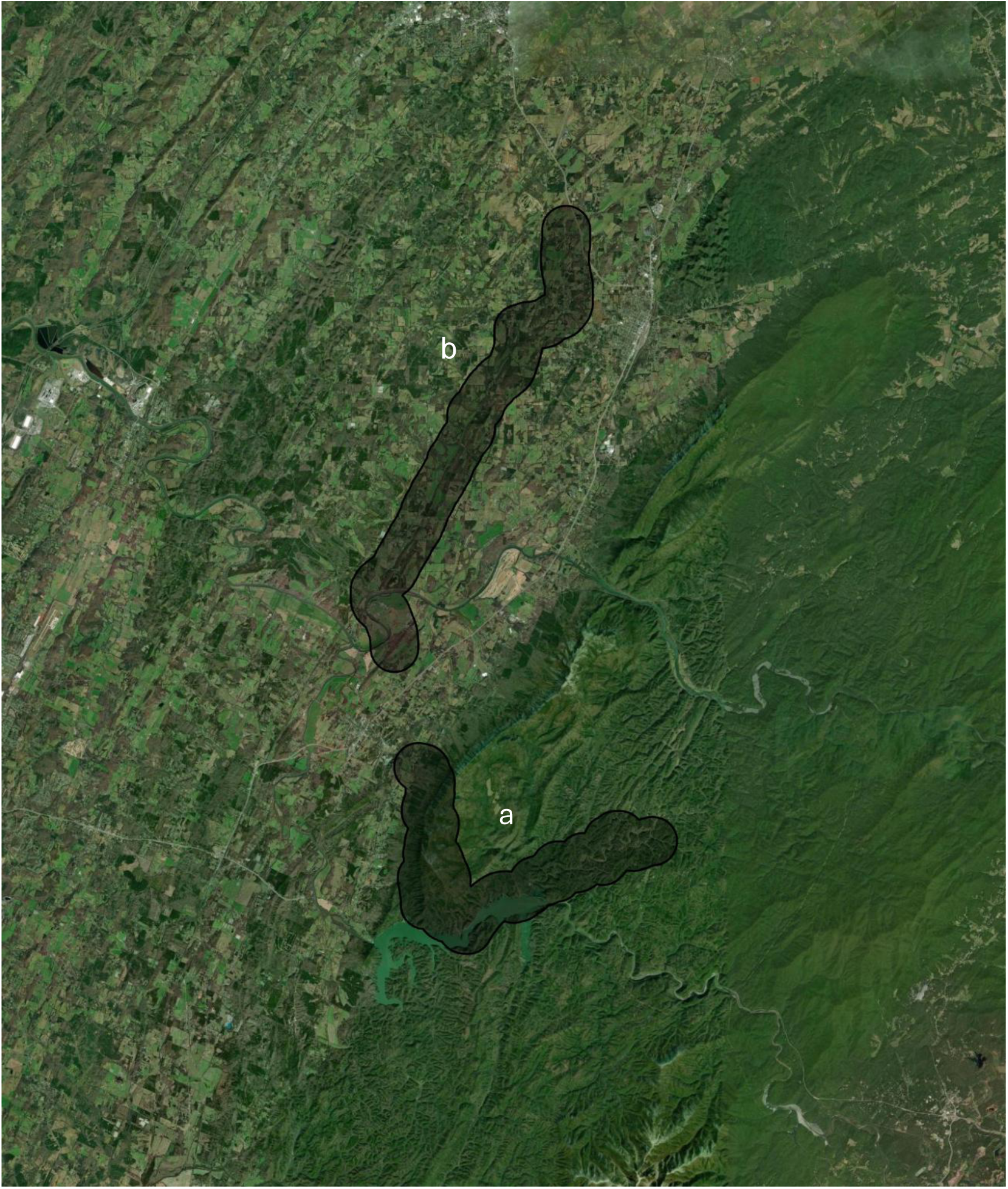
ArcGIS image of our forest (a) and farmland (b) transects with 1 km buffers

Our farmland transect was 23.8 kilometers long with patches of woodland surrounded by small farm fields. There was a single river crossing near the start of the transect, with no other detectable surface water aside from occasional field drainage ditches. We used ArcGIS to calculate percent canopy coverage within a 1-kilometer buffer. (Fig. 1).

### Mobile acoustic sampling methods

We then drove along each of these transects with an ultrasonic recorder (Titley Scientific Chorus) mounted on top of a vehicle. The roof mounting method is an effective way to obtain many recordings in a single transect route in mobile bat detection surveys (Whitby *et al*. 2014). Our recorder was fitted with an ultrasonic microphone, which allowed it to record the calls of foraging bats. We used bioacoustic analysis software (Kaleidoscope Pro 5.6.6, Wildlife Acoustics, Maynard, MA, U.S.A.) to identify bats based on these recorded calls. When using transects to estimate abundance, animal movement independent of the observer can cause bias; if animals are moving more quickly than the observer it can lead to inaccurately high estimations of abundance (Glennie et al. 2015). Thus, speed was maintained between 24.1 to 40.2 km/h along the transect routes. This speed is optimal to outpace flying foraging bats, greatly reducing the risk of recording the same individual multiple times. Recording began 10-15 minutes before sundown each night (D’Acunto *et al*. 2018). To control for any effect of time of night on which species are present along the transects, the order of transect sampling was alternated each night of sampling. In summer 2024, We collected 15 full nights of sampling (both farmland and forest were sampled) and two half nights (one where just the forest was sampled and another where only the farmland was sampled) yielding a final count of 16 transect runs for both habitats.

### Spectrogram analysis and species identification

When hunting, bats emit ultrasonic sound waves to echolocate their prey. These foraging calls differ in frequency, length, and change in frequency depending on the species of bat, making it possible to identify bat species from recordings of their calls. Among the species we detected, call frequencies typically range from 20 – 50+ kHz (NABat, North American Bat Monitoring Program United States, Canada 2025). We classified calls to species by using Kaleidoscope’s North American bat automatic identification feature and then checking spectrograms to finalize classification to species (Fig. 2).

**Fig. 2.**
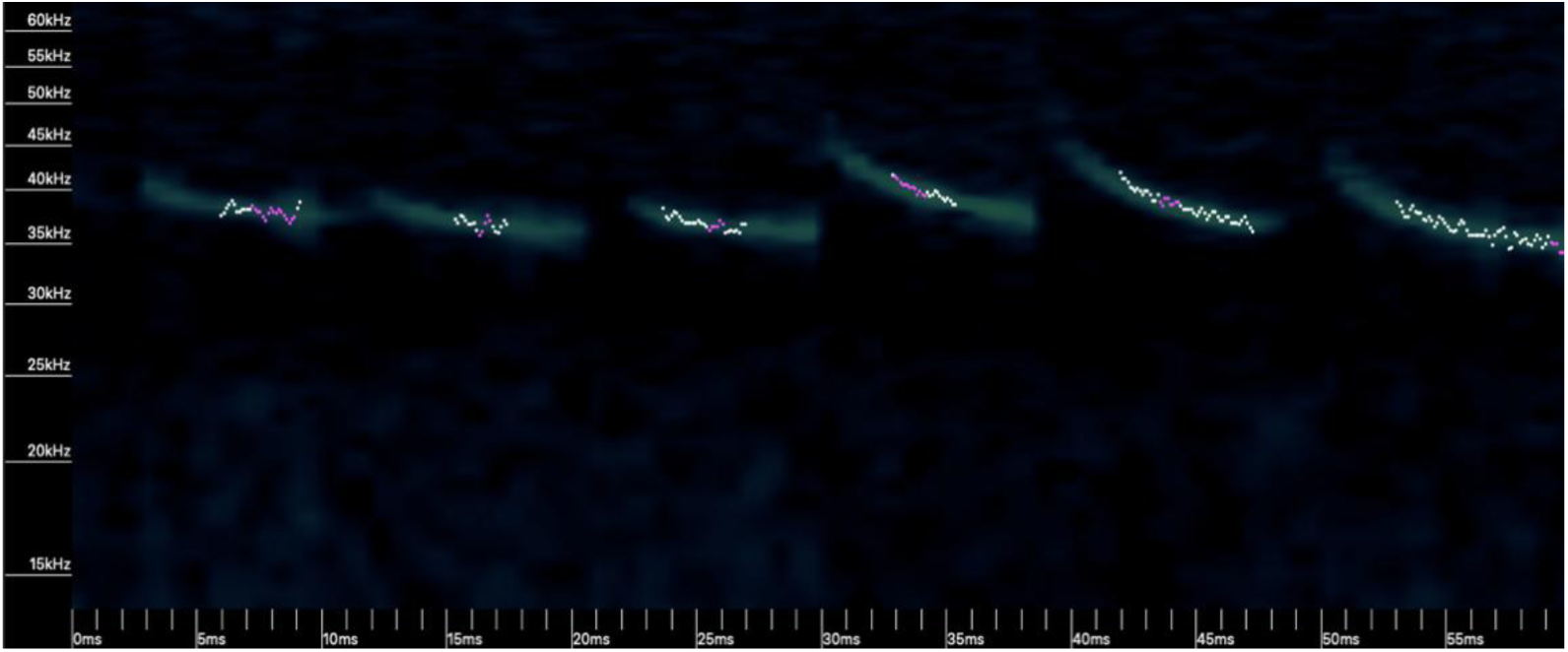
Example of spectrogram depicting the characteristic call signature (in kHz and ms) of *L. seminolus*

While Kaleidoscope’s Auto-ID detected many individual bat calls, it classified only 32.5% of them to species. We manually identified as many of the remaining 67.5% of recordings as possible by using a bat call characteristic guide published by Humboldt State University (HSU Bat Lab 2011) and comparing our spectrograms to known call characteristics of bat species whose ranges include our study site according to NABat (2025). We managed to identify all but 97 calls from the forest and farmland. These remaining recordings were not clear enough to identify any species, either due to overlap from multiple bats in the recording, other ultrasonic noise sources blocking out bat calls, or too few vocalizations within a given recording to reliably measure identification parameters. We were thus able to identify 89.4% of all bats encountered.

### Statistical analysis

We calculated the Shannon diversity index (Shannon 1948) for the farmland, the forest, and each individual section of the forest transect. Hutcheson’s t-test was used to test whether the two communities differed in diversity. Following this, we calculated post hoc exact binomial tests to identify between which species are most strongly contributing to this difference. We then calculated *β_Shannon_* (Jost 2007) to determine if habitat type (farmland versus forest) contributed to diversity versus each habitat being equivalent in species composition. *β_Shannon_* was also applied to our comparison of diversity between forest subtransects A, B, and C. Pairwise Bray-Curtis Dissimilarity scores (Bray and Curtis 1957) between the forest, its sections, and the farmland were calculated to describe the difference in community composition between the two sites, and between the forest subtransects. Data was visualized using the ggplot2 package v3.3.3. (Wickham 2016)

## Results

### Forest versus farmland

Our final sample size included 914 bats across both habitats, 583 in the forest and 331 in the farmland. Mean abundance of bats across species differed between forest and farmland (paired t-test, N = 9, t = 3.548856, P = 0.007522). We also compared the mean canopy coverage per square kilometer in ArcGIS which yielded 79.5% for the forest and 31.5% for the farmland. The same nine species were detected in both habitats (Fig. 3). However, eight of the nine species detected were more abundant in the intact forest than in the fragmented habitat of the farmland, with Seminole bats (*L. seminolus*), being the sole exception, being detected slightly more often in the farmland (Fig. 3). Due to species-specific responses, the relative abundance of species in forest versus farmland was significantly different overall (Fisher’s exact test, P= 0.0004998).

**Table 1:** *Post-hoc* exact binomial tests for each species comparing proportion of total detections that occurred in forest versus farmland to the expected ratio based on overall bat abundance (583:331). *α* = 0.0055 after Bonferroni correction.

| Species | N (detections across forest and farmland) | P |
| --- | --- | --- |
| Little brown (*Myotis lucifugus*) | 73 | 0.001393* |
| Big brown (*Eptesicus fuscus*) | 35 | 0.725463 |
| Eastern red (*Lasiurus borealis*) | 242 | 0.688290 |
| Hoary (*Lasiurus cinereus*) | 36 | 0.603491 |
| Evening (*Nycticeus humeralis*) | 125 | 0.019898 |
| Tricolored (*Perimyotis subflavis*) | 79 | 0.159578 |
| Silver haired (*Lasiurus noctivagans*) | 74 | 0.183699 |
| Seminole (*Lasiurus seminolus*) | 239 | 2.98E-06* |
| Northern (*Myotis septentrionalis*) | 11 | 0.347612 |

**Fig. 3.**
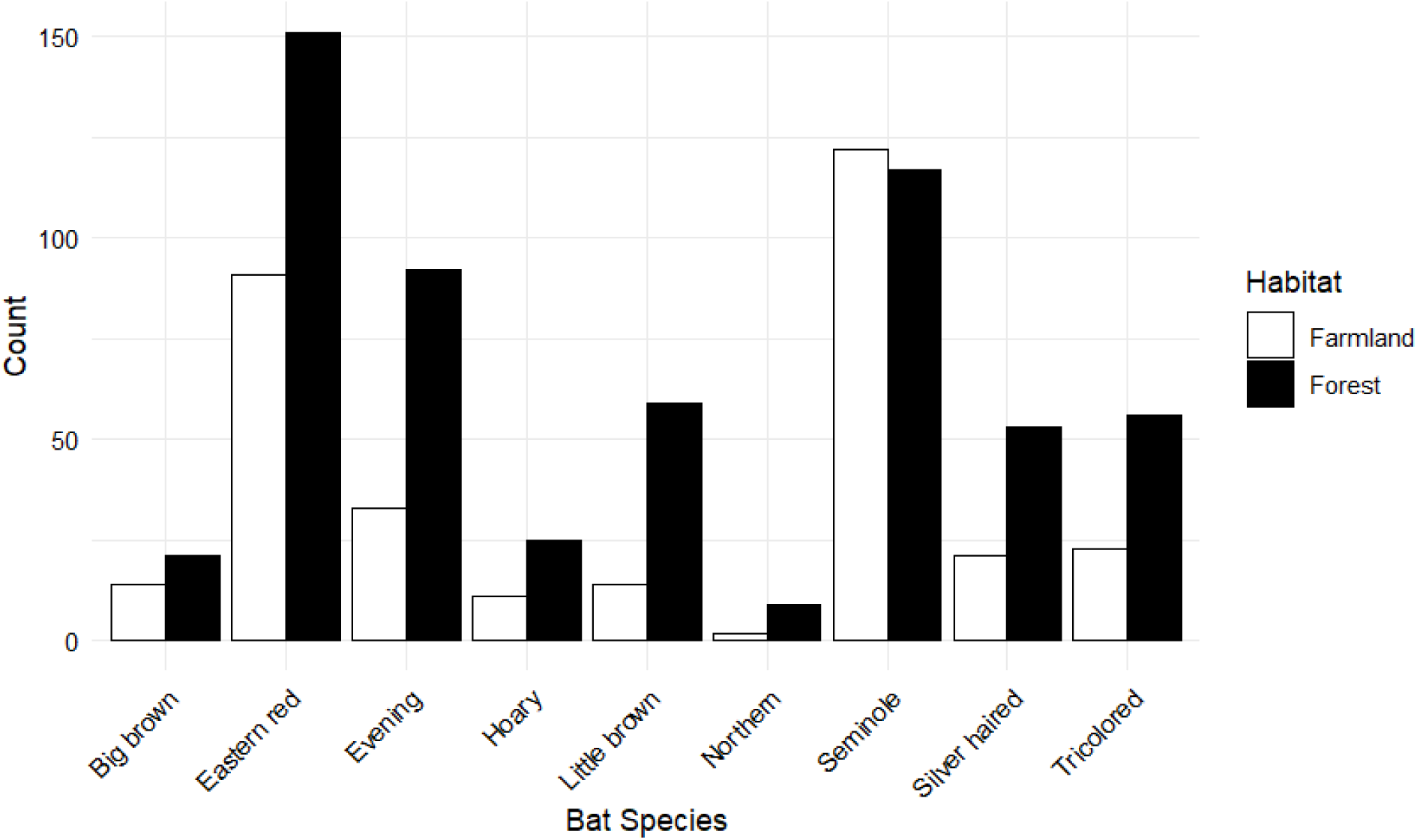
The abundance of each species in the farmland (white) and forest (black) there were more individuals of every species in the forest with the exception of Seminole bats.

### Diversity indices comparing forest and farmland

We also calculated the Shannon diversity index (H’), *β_Shannon_*, and Bray-Curtis dissimilarity between the forest and the farmland, (H’_forest_ = 1.96; H’_famland =_ 1.72; βShannon = 1.05; Bray-Curtis = 0.287) Together, these indices indicate that the forest community is more diverse than the fragmented farmland community, as confirmed by a Hutcheson’s t-test (t = 4.297, P = 0.0000205).

### Analysis of surface water’s effect on forest bat communities

We compared forest transect segments A, B, and C to determine if there were any significant differences due to the varying bodies of water along the transect. The subtransects had 241, 65, and 149 individuals in forest A, B, and C, respectively. Roughly 13.7 kilometers of the 25.75-kilometer transect was comprised of section A, just 3.2 kilometers were of section B, and the remaining 8.8 kilometers were section C. Abundance did not differ from the expected counts expected given subtransect length and total detections in the forest transect (Χ^2^ = 1.514 P = 0.469). Community composition differed among the three forest segments (Fisher’s exact test, N=306, P=0.0005). Post hoc pairwise Fisher’s exact tests showed that sections A and C significantly differed in species composition, (N=390, P=0.0005). However, no significant difference in community composition was found between forest subtransects A and B (N=306, P=0.2034). Sections B and C also did not significantly differ (N=214, P=0.0660), however, the sample was over 30% smaller than section A and C, reducing our achieved power and our ability to detect an effect of the same size that existed between sections A and B.

We calculated β_Shannon_ and pairwise Bray-Curtis dissimilarities between forest transect segments A, B, and C. (*β_Shannon_* 1.075) Our pairwise β_Shannon_ comparisons all show a moderate effect of habitat type on diversity.

Our β_Shannon_ indicates that habitat does contribute to overall diversity, especially when comparing A and C, *β_Shannon_* does find a larger difference when comparing A-B and B-C. At the start of sampling, some forest data was collected and not included in Table II due to the separation between forest subtransects A, B, and C not being established yet.

## Discussion

In the present study, we found that vesper bat abundance dropped by over 43% in the fragmented habitat of the farmland relative to the intact forest.

### Fragmentation effects due to edge effects and loss of habitat

When we compare the amount of forest canopy between our two habitats, we see a major difference, with 79.5% versus 32.5% coverage in the forest and farmland respectively. Further, forest patches in farmland have a greater proportion of edge to core habitat, and edge habitat tends to support less diversity due to changes in forest canopy structure (Blanchard et al., 2023) With such a major loss of habitat a decrease in community diversity in the farmland is expected, especially in the case of less-mobile species (Lu, *et al* 2012; Püttker et al 2013). However, vesper bats are highly mobile and should be able to cross significant areas of matrix between fragments when foraging. Despite vesper bats’ high mobility, we observed a drastic decrease in abundance in all but one identified species, Seminole bats. This suggests that as habitat fragmentation reduces overall canopy coverage drastic reductions in species abundance, regardless of dispersal ability, will follow. The number of Seminole bats was similar between the two habitats. When we compare the proportion of each species in the farmland community to those in the forest, only Seminole and little brown bats significantly changed their relative abundance—increasing relative to the other species. This relative abundance increase reflects a less severe decrease in absolute abundance for little brown bats and perhaps no decrease in abundance for Seminole bats. All other species comprised a similar proportion of their respective community, suggesting that seven of our nine species are decreasing in abundance at a similar rate. If we consider the species whose abundance significantly differed (that is, all but Seminole bats), we can see a clear correlation between the magnitude of canopy fragmentation and vesper bat abundance decrease.

### Canopy fragmentation effects on bat communities

Gorresen and Willig (2004) found that phyllostomid bat species richness peaks at intermediate levels of forest fragmentation. Further, Laforge et al. (2022) found that bat responses to forest cover on a landscape scale are species-specific, depending on preferences in roosting and foraging habitat; bats who rely on closed-canopy forests to roost and forage were positively associated with forest cover, whereas species such as little brown bats, that are known to forage in open habitat, were negatively affected. Our finding that the farmland bat community is less diverse than the forest’s is expected given the high degree of fragmentation and habitat loss. Absolute abundance was reduced by a similar proportion in most species, resulting in similar community composition. However, our post hoc exact binomial tests indicated that Seminole and little brown bats comprise a significantly greater proportion of the farmland community because their abundance did not decrease as much as the other species. Taken together these facts suggest that the observed species may be heavily reliant on canopy coverage within their habitat, with the exception of the Seminole bat. Extinction of core-species populations have been observed as a result of habitat fragmentation via habitat loss, negative edge effects, or both (Ewers and Didham 2007).

### Species-specific responses to habitat fragmentation

The bat community in the Cherokee National Forest is more diverse due to higher species evenness, although richness is identical in both communities. Some species, such as the Seminole bat, appear to be tolerant of canopy fragmentation, as they maintained similar abundance after fragmentation. The drop in abundance of other bat species suggests Seminole bats will have reduced interspecific competition and that they may be better adapted to small forest patches within the farmland matrix than other bat species. Hein, Castleberry, and Miller (2008) found that Seminole bats prefer forest corridor habitat for roosting in managed landscapes, finding that landscape features were more important than tree and plot-level characteristics when selecting roosting sites. Thus, Seminole bats could adapt to fragmented habitats and exploit them for roosting, foraging, or both, thereby reducing interspecific competition and making them relative winners in the context of canopy fragmentation (Fahrig *et al*. 2019). A study in central Chile found that insectivorous bat activity was higher at the edges created by human modification; however, they also found that some species such as cinnamon red bats (*Lasiurus varius*) thrive in smaller core habitat patches within fragmented matrices, indicating that they had adapted to successfully roost and forage within human modified habitat (Rodríguez-San and Simonetti 2013). Understanding the behavioral differences between species within an anthropogenic matrix will be essential to understanding species dynamics as human development inevitably further fragments forest ecosystems across the world (Rybicki and Haskin 2013).

### Mixed responses to fragmentation

It is evident that Seminole bats can thrive in a fragmented landscape while others struggle to adapt to the change in habitat composition (Rybicki, Abrego, and Ovaskainen 2020). Atkinson and Shorrocks (1981) found that dividing habitat into smaller pieces can foster the coexistence of multiple competitive species by creating aggregates of superior competitors. However, if this elimination or reduction of interspecific and intraspecific competition was the cause, we might expect other species to also maintain high abundance. Instead, only Seminole bats occurred in similar abundance in the farmland versus the forest.

Another hypothesis that could explain the varied responses to habitat fragmentation is differing predator-prey dynamics between bat species. Differing dietary preferences among species may influence which species are drawn to forage in fragmented habitats. Every species observed in our study is insectivorous, we often observed bats divebomb in efforts to catch flying insects as acoustic sampling occurred this summer. Rodríguez-San and Simonetti (2015) found that fragmented habitat with native forest patches within low-contrast matrices, such as the farmland examined here, could well support foraging insectivorous bat communities. Marcantonio *et al*. (2023) found that *Limenitis camilla* butterflies spent more time moving within a fragmented forest habitat than their well-connected habitat counterparts. We hypothesize that if insects within the farmland are responding like those in Marcantonio *et al*. (2023) with increased movement, it is possible that certain species such as Seminole bats could successfully forage in the fragmented habitat due to an increased amount of vulnerable flying insects. Studying insect behavior in response to habitat fragmentation could help us better understand bat community dynamics in fragmented habitats. Furthermore, in addition to roosting preferences, greater understanding of the dietary preferences of the local bat community could also provide insight into how each bat species may respond as habitat fragmentation continues and affects prey behavior and populations.

### Comparison of surface water effects on forest bat communities

The farmland transect does not intersect or run along flowing streams for most of its length, while the forest transect has several kilometers that run along streams. It is possible that this difference in surface water is affecting the difference in bat community composition and possibly bat abundance between forest and farmland. However, this possible surface water effect in addition to fragmentation effects is unlikely to explain the differences we saw.

We saw greater Bray-Curtis dissimilarity between our forest subtransects than between the forest and farmland. However, Bray-Curtis dissimilarity relies on the proportion of the community each species comprises and is susceptible to over or underestimation of dissimilarity when applied to a small sample size. Reduced sample size, and differences in sample size, as in our three forest subtransects (especially section B), may increase measures of dissimilarity relative to the dissimilarity seen between our two entire transects (Schmera and Eros 2006). The length difference between our forest subtransects could confound interpretation of the effect of water along the transect as the forest B (riparian) section is by far the shortest section of the transect, at just around 3.2 kilometers of the 25.75 kilometer transect. The comparison between the two other sections, forest A (dry) versus forest C (small streams), may provide a better indicator of the effect of water on bat foraging behavior and abundance. However, the observed magnitude of dissimilarity could still be exaggerated relative to the forest versus farmland comparison due to small sample sizes in these subtransects. While A and C are similar in length, there is a smaller difference in amount of surface water in both sections than when compared to B, which also has a much smaller sample size due to path of the available roads. The difference in relative abundance (community composition) across our three forest subtransects was significant according to our Fisher’s exact test; however, the only pairwise comparison that was statistically significant was the comparison of subtransects B and C, which have differences in both surface water and in length, with subtransect C being 2.75 times as long as B. Hutcheson’s t-test revealed a statistically significant difference in diversity only between subtransects B and C with or without Bonferroni correction (Table 2); Together, these results suggest that the different amount of surface water between the forest and farmland is probably not a strong confounding variable but is worthy of further study. Given the fact that the riparian (B) forest section was so much shorter in length, future research could elucidate whether bat communities are affected by amount of surface water by sampling transects of equal lengths and differing amounts of surface water when comparing fragmented and unfragmented habitat.

**Table 2:** Pairwise comparison of bat diversity between forest transect segments.

| **Diversity indices:** | **A-B** | **A-C** | **B-C** |
| --- | --- | --- | --- |
| Bray-Curtis dissimilarity | 0.58 | 0.32 | 0.39 |
| $\beta_{Shannon}$ | 1.15 | 1.005 | 1.09 |
| Hutcheson's t-test (t) | 1.848426 | 1.502554 | 2.906370 |
| Hutcheson's t-test (P) | 0.067227 | 0.133798 | 0.004461 |

In conclusion there is a significant reduction of abundance in the fragmented habitat versus the intact forest. However, proportional differences in species composition between the forest and farmland were mostly insignificant. Seminole bats are a notable exception with similar abundance in both habitats, comprising a much higher proportion of the farmland community. Little brown bats also comprised a larger proportion of the farmland community than the forest community. With differing responses to fragmentation on a species-by-species basis, our data corroborates Fahrig (2003) and Rybicki and Haskin’s (2013) assertions that there can be a positive or negative response to habitat fragmentation. We observed a significant effect of fragmentation on the community composition of a highly mobile set of species, suggesting that an organism’s ability to move long distances may not be able to counteract the effects of habitat isolation from fragmentation. The overall drop in abundance of such a highly mobile group suggests that habitat loss is the driving factor more than fragmentation itself. On a related note, given how mobile vesper bats are, we cannot rule out the possibility of spillover of the forest community into the farmland during nightly foraging; or vice versa. However, such spillover would reduce the differences in abundance and community composition we observed, making our results a conservative estimate of the effects of habitat loss and fragmentation. Observing an area of fragmented forest with more distance from intact forest would elucidate whether spillover effects are significant. A better understanding of bat foraging ranges in these populations via tagging different species would also help determine whether spillover is a confounding factor. Seminole bat foraging behavior is of interest in this context given their apparent ability to maintain high abundance in a fragmented habitat.

On the same note, our understanding of bat community dynamics would benefit not only from the identification of foraging ranges of all the species in our dataset, but also by identifying the effect of fragmentation on the flying insects that local vesper bats are preying upon. If there were a strong correlation between increasing fragmentation and decreasing insect abundance, this could indicate a potential bottom-up effect that results in a loss of bat abundance. Thus, further research into bat foraging ranges and insect response to fragmentation would help elucidate which effects of habitat fragmentation are reducing insectivorous bat diversity. Our fragmented farmland transect consisted of small, highly divided, edge-heavy habitat patches within a large matrix, it is possible that negative effects of habitat fragmentation could be minimized by clustering habitat fragments, thereby reducing the loss of core habitat, and allowing more species to persist in wake of land sharing efforts (Rybicki and Haskin, 2013). Even though we cannot conclude that habitat fragmentation is the sole factor reducing bat biodiversity (e.g. insecticide use in farmlands may play a role), our findings indicate an adverse effect of habitat fragmentation on Eastern United States bat community composition and abundance.

## Acknowledgements

Thank you to the Ronald E. McNair Post-Baccalaureate Achievement Program Department of Education Grant award P217A220116 – 25, which funded this research.

## Statements and Declarations

### Author Contributions

All authors participated in conceptualizing and designing this study. Data collection was performed by Ethan J. Ramey. All authors contributed to data analysis and interpretation. All authors participated in drafting and revising this transcript. All authors have read and approved the submission of the final manuscript.

## References

1. Atkinson WD, Shorrocks B (1981) Competition on a divided and ephemeral resource: A simulation model. The Journal of Animal Ecology 50:461. 10.2307/4067

2. Beilke EA, O’Keefe JM (2023) Bats reduce insect density and defoliation in temperate forests: An exclusion experiment. Ecology 104:e3903. 10.1002/ecy.3903

3. Bernard E, Fenton MB (2007) Bats in a fragmented landscape: Species composition, diversity and habitat interactions in savannas of Santarém, Central Amazonia, Brazil. Biological Conservation 134:332–343. 10.1016/j.biocon.2006.07.021

4. Blanchard G, Barbier N, Vieilledent G, et al (2023) UAV-Lidar reveals that canopy structure mediates the influence of edge effects on forest diversity, function and microclimate. Journal of Ecology 111:1411–1427. 10.1111/1365-2745.14105

5. Bray JR, Curtis JT (1957) An ordination of the upland forest communities of southern wisconsin. Ecological Monographs 27:325–349. 10.2307/1942268

6. Chetcuti J, Kunin WE, Bullock JM (2020) Habitat fragmentation increases overall richness, but not of habitat-dependent species. Frontiers In Ecology Evol 8:607619. 10.3389/fevo.2020.607619

7. Cueva Salcedo HDL, Fenton MB, Hickey MBC, Blake RW (1995) Energetic consequences of flight speeds of foraging red and hoary bats (L*asiurus borealis* and L*asiurus cinereus* ; Chiroptera: *Vespertilionidae)*. Journal of Experimental Biology 198:2245–2251. 10.1242/jeb.198.11.2245

8. D’Acunto LE, Pauli BP, Moy M, et al (2018) Timing and technique impact the effectiveness of road-based, mobile acoustic surveys of bats. Ecology and Evolution 8:3152–3160. 10.1002/ece3.3808

9. Dechen Quinn AC, Williams DM, Porter WF (2013) Landscape structure influences space use by white-tailed deer. Journal of Mammal 94:398–407. 10.1644/11-MAMM-A-221.1

10. Dunning JB, Danielson BJ, Pulliam HR (1992) Ecological processes that affect populations in complex landscapes. Oikos 65:169. 10.2307/3544901

11. Emlen ST (1972) An experimental analysis of the parameters of bird song eliciting species recognition. Behavior 41:130–171. 10.1163/156853972X00248

12. Ewers RM, Didham RK (2007) The effect of fragment shape and species’ sensitivity to habitat edges on animal population size. Conservation Biology 21:926–936. 10.1111/j.1523-1739.2007.00720.x

13. Fahrig L (2017) Ecological responses to habitat fragmentation per se. Annual Review of Ecology, Evolution, and Systematics 48:1–23. 10.1146/annurev-ecolsys-110316-022612

14. Fahrig L (2002) Effect of habitat fragmentation on the extinction threshold: a synthesis. Ecological Applications 12:346–353. 10.1890/1051-0761(2002)012%255B0346:EOHFOT%255D2.0.CO;2

15. Fahrig L (2003) Effects of habitat fragmentation on biodiversity. Annual Review of Ecology, Evolution, and Systematics 34:487–515. 10.1146/annurev.ecolsys.34.011802.132419

16. Fahrig L (2007) Non-optimal animal movement in human-altered landscapes. Functional Ecology 21:1003–1015. 10.1111/j.1365-2435.2007.01326.x

17. Fahrig L, Arroyo-Rodríguez V, Bennett JR, et al (2019) Is habitat fragmentation bad for biodiversity? Biological Conservation 230:179–186. 10.1016/j.biocon.2018.12.026

18. Federico P, Hallam TG, McCracken GF, et al (2008) Brazilian free-tailed bats as insect pest regulators in transgenic and conventional cotton crops. Ecological Applications 18:826–837. 10.1890/07-0556.1

19. Fenton MB, Bell GP (1981) Recognition of species of insectivorous bats by their echolocation calls. Journal of Mammalogy 62:233–243. 10.2307/1380701

20. Fletcher RJ (2005) Multiple edge effects and their implications in fragmented landscapes. Journal of Animal Ecology 74:342–352. 10.1111/j.1365-2656.2005.00930.x

21. Fletcher RJ, Didham RK, Banks-Leite C, et al (2018) Is habitat fragmentation good for biodiversity? Biological Conservation 226:9–15. 10.1016/j.biocon.2018.07.022

22. Frick WF, Kingston T, Flanders J (2020) A review of the major threats and challenges to global bat conservation. Annals of the New York Academy of Sciences 1469:5–25. 10.1111/nyas.14045

23. Fullard JH, Dawson JW (1997) The echolocation calls of the spotted bat E*uderma maculatum* are relatively inaudible to moths. Journal of Experimental Biology 200:129–137. 10.1242/jeb.200.1.129

24. Glennie R, Buckland ST, Thomas L (2015) The effect of animal movement on line transect estimates of abundance. PLoS One 10:e0121333. 10.1371/journal.pone.0121333

25. Gorresen PM, Willig MR (2004) Landscape responses of bats to habitat fragmentation in Atlantic forest of Paraguay. Journal of Mammalogy 85:688–697. 10.1644/BWG-125

26. Haddad NM, Brudvig LA, Clobert J, et al (2015) Habitat fragmentation and its lasting impact on Earth’s ecosystems. Science Advances 1:e1500052. 10.1126/sciadv.1500052

27. Hanski I (2011) Habitat loss, the dynamics of biodiversity, and a perspective on conservation. AMBIO 40:248–255. 10.1007/s13280-011-0147-3

28. Heide K, Friesen L, Martin V, et al (2023) Before-and-after evidence that urbanization contributes to the decline of a migratory songbird. Avian Conservation and Ecology 18:art15. 10.5751/ACE-02366-180115

29. Hein CD, Castleberry SB, Miller KV (2008) Sex-specific summer roost-site selection by seminole bats in response to landscape-level forest management. J Mammal 89:964–972. 10.1644/07-MAMM-A-335.1

30. Henle K, Davies KF, Kleyer M, et al (2004) Predictors of species sensitivity to fragmentation. Biodiversity and Conservation 13:207–251. 10.1023/B:BIOC.0000004319.91643.9e

31. Hoyt JR, Kilpatrick AM, Langwig KE (2021) Ecology and impacts of white-nose syndrome on bats. Nature Reviews Microbiology 19:196–210. 10.1038/s41579-020-00493-5

32. Jones G, Holderied MW (2007) Bat echolocation calls: adaptation and convergent evolution. Proceding of the Royal Socity B 274:905–912. 10.1098/rspb.2006.0200

33. Jost L (2007) Partitioning diversity into independent alpha and beta components. Ecology 88:2427–2439. 10.1890/06-1736.1

34. Krauss J, Bommarco R, Guardiola M, et al (2010) Habitat fragmentation causes immediate and time-delayed biodiversity loss at different trophic levels. Ecology Letters 13:597–605. 10.1111/j.1461-0248.2010.01457.x

35. Kupfer JA, Malanson GP, Franklin SB (2006) Not seeing the ocean for the islands: the mediating influence of matrix-based processes on forest fragmentation effects. Global Ecology and Biogeography 15:8–20. 10.1111/j.1466-822X.2006.00204.x

36. Laforge A, Barbaro L, Bas Y, et al (2022) Road density and forest fragmentation shape bat communities in temperate mosaic landscapes. Landscape and Urban Planning 221:104353. 10.1016/j.landurbplan.2022.104353

37. Law BS, Dickman CR (1998) The use of habitat mosaics by terrestrial vertebrate fauna: implications for conservation and management. Biodiversity and Conservation 7:323–333. 10.1023/A:1008877611726

38. Lu N, Jia C-X, Lloyd H, Sun Y-H (2012) Species-specific habitat fragmentation assessment, considering the ecological niche requirements and dispersal capability. Biological Conservation 152:102–109. 10.1016/j.biocon.2012.04.004

39. Marcantonio M, Voda R, Da Re D, et al (2023) The effect of habitat on insect movements: experimental evidence from wild-caught butterflies. Insects 14:737. 10.3390/insects14090737

40. Meyer CFJ, Struebig MJ, Willig MR (2016) Responses of tropical bats to habitat fragmentation, logging, and deforestation. In: Voigt CC, Kingston T (eds) Bats in the Anthropocene: Conservation of Bats in a Changing World. Springer International Publishing, Cham. 10.1007/978-3-319-25220-9_4

41. Moreno CE, Halffter G (2000) Assessing the completeness of bat biodiversity inventories using species accumulation curves. Journal of Applied Ecology 37:149–158. 10.1046/j.1365-2664.2000.00483.X

42. North American Bat Monitoring Program | United States | Canada. In: NABat. https://www.nabatmonitoring.org. Accessed 2 Apr 2025

43. Pardini R, bueno ADA, gardner TA, et al (2010) beyond the fragmentation threshold hypothesis: regime shifts in biodiversity across fragmented landscapes. PLoS One 5:e13666. 10.1371/journal.pone.0013666

44. Park KJ (2015) Mitigating the impacts of agriculture on biodiversity: bats and their potential role as bioindicators. Mammalian Biology 80:191–204. 10.1016/j.mambio.2014.10.004

45. Parker TH, Stansberry BM, Becker CD, Gipson PS (2005) Edge and area effects on the occurrence of migrant forest songbirds. Conservation Biology 19:1157–1167. 10.1111/j.1523-1739.2005.00107.x

46. Pedro AR-S, Simonetti JA (2013) Foraging activity by bats in a fragmented landscape dominated by exotic pine plantations in central chile. Acta Chiropterologica 15:393–398. 10.3161/150811013X679017

47. Püttker T, Bueno AA, Dos Santos De Barros C, et al (2013) Habitat specialization interacts with habitat amount to determine dispersal success of rodents in fragmented landscapes. Journal of Mammalogy 94:714–726. 10.1644/12-MAMM-A-119.1

48. Rocha R, Ovaskainen O, López-Baucells A, et al (2018) Secondary forest regeneration benefits old-growth specialist bats in a fragmented tropical landscape. Scientific Reports 8:3819. 10.1038/s41598-018-21999-2

49. Rodríguez-San Pedro A, Simonetti JA (2015) The relative influence of forest loss and fragmentation on insectivorous bats: does the type of matrix matter? Landscape Ecology 30:1561–1572. 10.1007/s10980-015-0213-5

50. Rybicki J, Abrego N, Ovaskainen O (2020) Habitat fragmentation and species diversity in competitive communities. Ecology Letters 23:506–517. 10.1111/ele.13450

51. Rybicki J, Hanski I (2013) Species–area relationships and extinctions caused by habitat loss and fragmentation. Ecology Letters 16:27–38. 10.1111/ele.12065

52. Schmera D, Eros T (2006) Estimating sample representativeness in a survey of stream caddisfly fauna. International Journal of Limnology 42:181–187. 10.1051/limn/2006019

53. Shannon CE (1948) A mathematical theory of communication. Bell System Technical Journal 27:379–423. 10.1002/j.1538-7305.1948.tb01338.x

54. Szitár K, Tölgyesi C, Deák B, et al (2023) Connectivity and fragment size drive plant dispersal and persistence traits in forest steppe fragments. Frontiers in Ecology and Evolution 11:1155885. 10.3389/fevo.2023.1155885

55. Tanner JE (2005) Edge effects on fauna in fragmented seagrass meadows. Austral Ecology 30:210–218. 10.1111/j.1442-9993.2005.01438.x

56. Tscharntke T, Steffan-Dewenter I, Kruess A, Thies C (2002) Contribution of small habitat fragments to conservation of insect communities of grassland–cropland landscapes. Ecological Applications 12:354–363. 10.1890/1051-0761(2002)012%255B0354:COSHFT%255D2.0.CO;2

57. Whitby MD, Carter TC, Britzke ER, Bergeson SM (2014) Evaluation of mobile acoustic techniques for bat population monitoring. Acta Chiropterologica 16:223–230. 10.3161/150811014X683417

58. White AM, Holland TG, Abelson ES, et al (2022) Simulating wildlife habitat dynamics over the next century to help inform best management strategies for biodiversity in the Lake Tahoe Basin, California. Ecology and Society 27:art31. 10.5751/ES-13301-270231

59. Wickham H. Ggplot2: Elegant graphics for data analysis. 2nd ed. Cham, Switzerland: Springer International Publishing; 2016. 260 p.

60. Wordley CFR, Sankaran M, Mudappa D, Altringham JD (2018) Heard but not seen: Comparing bat assemblages and study methods in a mosaic landscape in the Western Ghats of India. Ecology and Evolution 8:3883–3894. 10.1002/ece3.3942

61. Yuan R, Zhang N, Zhang Q (2024) The impact of habitat loss and fragmentation on biodiversity in global protected areas. Science of the Total Environment 931:173004. 10.1016/j.scitotenv.2024.173004

